# PCA-based spatial domain identification with state-of-the-art performance

**DOI:** 10.1101/2024.07.29.605550

**Authors:** Darius P. Schaub, Behnam Yousefi, Nico Kaiser, Robin Khatri, Victor G. Puelles, Christian F. Krebs, Ulf Panzer, Stefan Bonn

## Abstract

The identification of biologically meaningful domains is a central step in the analysis of spatial transcriptomic data. Following Occam’s razor, we show that a simple PCA-based algorithm for spatial domain identification rivals the performance of ten competing state-of-the-art methods across six single-cell spatial transcriptomic datasets. Our reductionist approach, NichePCA, provides researchers with intuitive domain interpretation and excels in execution speed, robustness, and scalability.

## Main

Single-cell spatial transcriptomics (ST) extends our understanding beyond single-cell sequencing by revealing information about both individual cells and cell collectives. Similar to categorizing distinct cell types, these collectives can be classified into functionally or phenotypically distinct groups, known as spatial domains or niches. This allows us to describe disease pathology and dynamics in terms of cell-cell interactions and domain-specific cell-type distributions. Accurately identifying spatial domains is thus crucial to leverage the full potential of ST data. In recent years many sophisticated methods have been developed to identify tissue domains from ST data in an unsupervised manner^1–3^, spanning from graph neural network-based^4–10^ to Bayesian methods^11^. These methods generally follow the same sequence of high-level steps: beginning with graph construction, then neighborhood embedding, and finally clustering. We call this the neighborhood embedding paradigm. Within this paradigm, most methods try to achieve superior domain identification performance by introducing complexities in one or more of the underlying substeps e.g., by using graph neural networks as non-linear neighborhood embedding functions.

In this work, we investigate how much complexity is necessary for a model to achieve state-of-the-art spatial domain identification performance. Following Occam’s razor, we present a simple PCA-based workflow, NichePCA, and compare its domain identification performance to competing, more complex methods. The algorithm relies on only four steps: First, a k-nearest-neighbor graph is built using the segmented spatial data, with nodes representing cells and edges representing spatial cell adjacency. Second, the gene expression is normalized and log-transformed per cell and the mean gene expression of each cell and its neighbors is calculated. Third, PCA is performed on the aggregated gene expression counts to obtain a low-dimensional representation. Fourth, the spatial domains are identified using Leiden^12^ clustering. Note that the only difference between NichePCA and the common single-cell transcriptomic workflow for cell type identification is the spatial aggregation per cellular neighborhood.

We first assessed the performance of NichePCA by following the recently published spatial domain identification benchmark of Yuan et al.^1^. In their study, 13 spatial clustering methods were evaluated on multiple real and simulated single-cell ST datasets, each with expert-validated ground truth annotations. We extended this list by two additional methods, SpatialPCA^13^ and Banksy^14^. We considered three of Yuan et al.’s real datasets (our Datasets 1-3) that were generated using MERFISH^15^, BaristaSeq^16^, and STARmap^17^ technologies (see Methods). The performance of identifying spatial domains was evaluated against the ground truth annotations using the Normalized Mutual Information (NMI), Homogeneity (HOM), and Completeness (COM) scores. Fig. 1a shows the spatial domain identification performance of different methods across Datasets 1-3. As shown, Banksy demonstrates the highest performance, followed closely by NichePCA. The identified spatial domains by different methods along with their ground truth annotation for a selected sample of Dataset 1 are shown in Fig. 1b. We found that NichePCA outperforms Banksy on Dataset 1 and 2 after optimizing the number of nearest neighbors per dataset (Extended Data Fig. 1). An ablation experiment further validated that NichePCA combines minimal complexity with maximal performance within its algorithmic framework (Extended Data Fig. 2).

**Figure 1:**
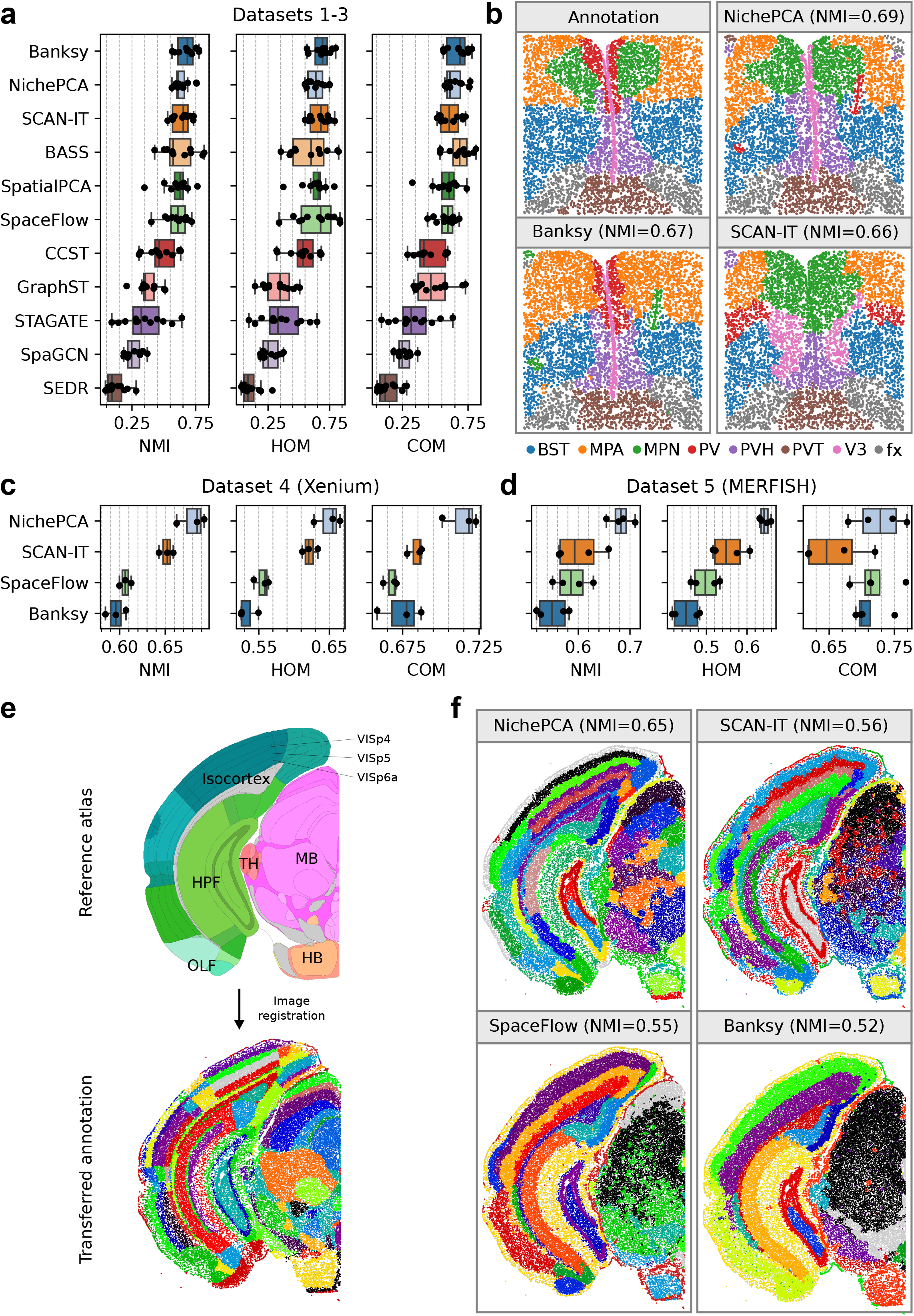
Quantitative performance evaluation on Datasets 1-5. **a**, Performance of different spatial domain identification methods on Datasets 1-3 in terms of NMI, HOM, and COM scores. The dots indicate the performance per sample. For SCAN-IT, BASS, SpaceFlow, CCST, GraphST, STAGATE, SpaGCN, and SEDR we adopted the results from Yuan et al.^1^. For NichePCA, Banksy, and SpatialPCA we followed an analogous benchmarking setup as Yuan et al. to ensure comparability (see Methods). **b**, Exemplary domain identification results for NichePCA, Banksy, and SCAN-IT on a selected sample of Dataset 1 (MERFISH) and corresponding ground truth annotations. In brackets, we provide the performance in terms of NMI for each method on this specific sample. **c, d**, Performance of NichePCA, SCAN-IT, SpaceFlow, and Banksy on Datasets 4 and 5, respectively, in terms of NMI, HOM, and COM scores. The dots show the performance per sample. We optimized each method on each dataset separately to ensure a fair comparison. **e**, (top) Schematic domain annotation taken from the Allen Brain Reference Atlas^27^ with labeled high-level areas. We additionally highlight some fine-grained domains inside the visual cortex. (bottom) Our ground truth domain annotations after image registration (see Methods) projected onto the cell positions within one half of a selected sample of Dataset 5 (MERFISH). We only show color codings but omit explicit cluster labels to preserve readability. Cells that could not be assigned to any domain have been removed. All metrics for this sample are calculated using the resulting fine-grained cluster assignments. **f**, Domain identification results for NichePCA, SCAN-IT, SpaceFlow, and Banksy on the same sample as in (e) with corresponding NMI scores in brackets. The NMI scores are calculated on both halves of the brain sample using the most fine-grained ground truth annotations.

While Datasets 1-3 contain less than 6,000 cells per sample, state-of-the-art spatial sequencing technology can feature more than 100,000 cells. This recent technological improvement, coupled with the increased performance of recent image registration and cell segmentation workflows, can significantly improve the quality and quantity of information that can be retrieved from ST data (see Methods). To understand the domain identification performance on state-of-the-art ST data, we next evaluated the four top-performing methods on two recent 10x Xenium (Dataset 4) and MERFISH (Dataset 5) brain datasets, using recent registration and segmentation technology (see Methods). In total, these datasets contain seven samples averaging about 100,000 cells per sample (Supplementary Table 1-2). Ground truth domain annotations for Dataset 4 were taken from Bhuva et al.^2^. For Dataset 5, we generated ground truth domain annotations by registering annotations from the Allen Brain Reference Atlas (see Methods). Since we were not able to execute BASS^11^ and SpatialPCA^13^ within a memory budget of 128 GB on any of the samples, we instead included SpaceFlow^6^, the next best algorithm, in this comparison. As shown in Fig. 1c-d, NichePCA significantly outperforms current state-of-the-art methods in all metrics on Datasets 4 and 5. Interestingly, Banksy, the best-performing method on Datasets 1 to 3, performs rather poorly on the more recent Datasets 4 and 5. On Dataset 4 NichePCA achieved a median NMI of 0.69, HOM of 0.65, and COM of 0.72, surpassing Banksy’s median NMI by 15, HOM by 25, and COM by 6 percentage points. For Dataset 5, NichePCA achieved a median NMI of 0.68, HOM of 0.64, and COM of 0.73, surpassing Banksy’s median NMI by 24, HOM by 42, and COM by 4 percentage points. We found this performance gap to persist when considering only transcripts inside the nuclei for Dataset 4, indicating that NichePCA’s superior performance is robust to changes in the cell segmentation procedure (Extended Data Fig. 3). The ground truth annotations and domain identification results of a selected sample from Dataset 5 are shown in Fig. 1e and 1f, respectively. NichePCA successfully identifies the separate layers in the cortex and even distinguishes different functional areas within these layers.

Many spatial domain identification algorithms do not support the integration of multiple samples. Recently, Yuan et al. combined SpaceFlow and Harmony^18^ to obtain coherent spatial domain clusters across multiple samples. NichePCA is naturally compatible with Harmony, as both rely on PCA, allowing for seamless multi-sample spatial domain identification. To assess this compatibility quantitatively we used the same MERFISH dataset^19^ as Yuan et al. (Dataset 6), containing 31 samples (see Methods). NichePCA outperforms the workflow developed by Yuan et al. both on a per-sample level as well as on all samples combined (Extended Data Fig. 4a, Fig. 2a). Three samples of identified domains along with the ground truth annotations are presented in Fig. 2b-d.

**Figure 2:**
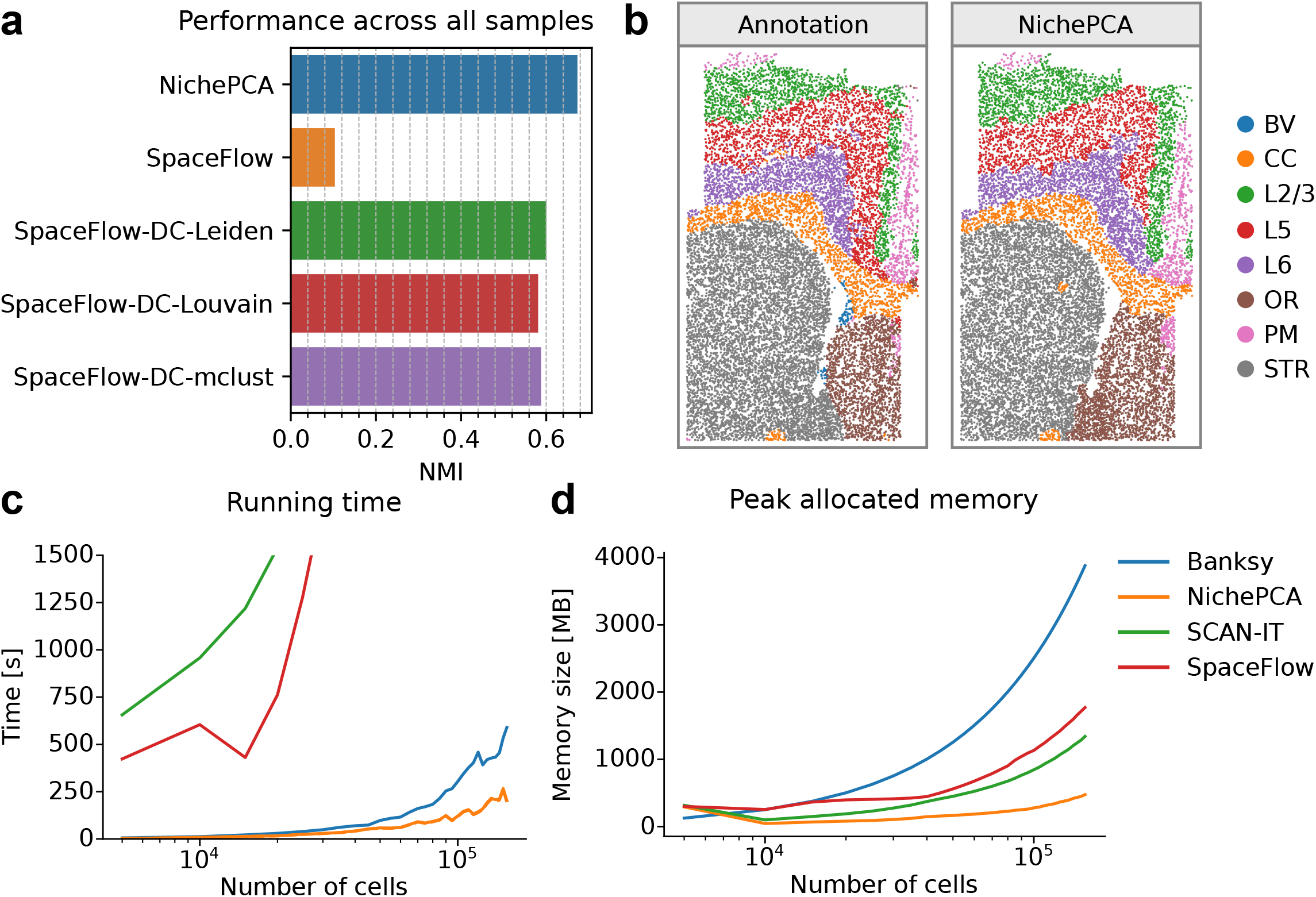
Multi-sample domain identification performance and scalability analysis. **a**, Spatial domain identification performance across all 31 samples of Dataset 6 in terms of NMI score for NichePCA, SpaceFlow, and different variants of SpaceFlow, that have been developed by Yuan et al.^1^. All results apart from the ones for NichePCA are adopted from Yuan et al.^1^. Since the NMI score is calculated across all samples, cluster labels for the same spatial domain must be preserved across samples to achieve high scores. **b**, Exemplary domain identification results for NichePCA on a selected sample of Dataset 6 and its corresponding ground truth annotations. Abbreviations for the different annotated domains are provided in the color legend. We manually matched the color coding of the NichePCA clusters to that of the ground truth domains. **c, d**, Peak allocated memory in megabytes (MB) and running time in seconds on subsampled versions (5,000-155,000 cells) of a sample from Dataset 4 for NichePCA, Banksy, SCAN-IT, and SpaceFlow. For a fair comparison, all methods are run solely on CPU (see Methods).

We further assessed the different workflows in terms of their running time and memory consumption (Fig. 2e and 2f). For this, we selected a sample of Dataset 4 and randomly subsampled it to different numbers of cells ranging from 5,000 to 155,000. Then, we applied Banksy, SCAN-IT, SpaceFlow, and NichePCA to each subsampled data. We found that NichePCA is by far the fastest and most memory-efficient method. The measured clustering times show that NichePCA primarily consumes time during the Leiden clustering step (Extended Data Fig. 4b-c). This could be further optimized by utilizing GPU implementations of the Leiden algorithm^20^.

NichePCA inherits the advantages of PCA, such as the interpretation of each principal component (PC) in relation to genes. We can assess the variance explained as well as the top genes contributing to the variation in the data. For instance, in the analysis of MERFISH brain data, the first PC explains approximately 40% of the total variance present in the data, and the top genes contributing to it include oligodendrocyte markers such as *Olig1* and *Sox8* (Extended Data Fig. 5a-b). Directly relating gene contribution to cluster definition may involve post-hoc analysis, such as differential gene expression between clusters. However, this does not explain how clusters relate to data structure and how they are defined in gene space. We can project clusters and their centroids to PCA space (Extended Data Fig. 5c). This allows us to identify the top genes contributing to each cluster by multiplying the PC coordinates of cluster centroids by the PCA feature loadings (see Methods). As an illustration, we examine cluster 29, which corresponds to the hippocampus region CA1 stratum radiatum (CA1sr) (Extended Data Fig. 5d-e). CA1 neurons are critical for hippocampal-dependent memory formation. *Sstr4*, a gene involved in the selection of memory strategies^21^, is the gene with the highest positive contribution to the cluster 29 centroid. *Gpr161*, a gene localized in the hippocampus CA1 region^22^, is also among the top four genes (Extended Data Fig. 5f). Among the genes with the highest negative contribution are those with unspecific expression in the CA1 region, such as *Olig1*, a marker for mature oligodendrocytes (Extended Data Fig. 5g).

In summary, this work introduces NichePCA, a simple neighborhood embedding-based algorithm for spatial domain identification. It matches or surpasses the performance of current state-of-the-art methods and offers significant scalability advantages. Moreover, NichePCA’s simplicity allows for the easy interpretation of clustering results. We believe NichePCA can serve as a quick and efficient first-stop solution for identifying spatial domains, similar to how cell-level PCA is used for cell-type identification. While we acknowledge that the space of possible methods following the neighborhood embedding paradigm is far from explored, our empirical results suggest that higher complexity within this paradigm does not necessarily lead to better performance.

## Methods

### NichePCA

First, we normalize the raw spatial transcriptomics data to sum to their medians and then apply a log-transformation, respectively, using the *normalize_total* and *log1p* functions of the Scanpy (v1.9.8) Python package^23,24^. Next, we create a spatial graph with nodes representing cells and edges representing spatial adjacency between cells. To define cell adjacencies, either a distance threshold or a specified number (K) of nearest neighbors (KNN) can be used. By default, we use the KNN approach. Importantly, we ensure the graph is undirected by adding missing edges to the directed ones. The resulting cell neighborhood graph also includes self-loops. Then, for each cell, we calculate the mean gene expression of its neighbors, including the cell itself, and perform PCA on these aggregated gene expression vectors. Finally, the spatial domains are identified using Leiden clustering. In total, NichePCA depends on three hyperparameters: the number of nearest neighbors K, the number of principal components, and the Leiden resolution.

### Existing spatial clustering methods

Here we provide a brief overview of existing spatial clustering methods, including Banksy^14^, SpatialPCA^13^, BASS^11^, and SCAN-IT^7^.

**Banksy** uses three different cell- and neighborhood-level representations, including the raw gene expression of the cell, the aggregated raw expression of the neighboring cells (based on KNN), and a gene expression gradient in the cellular neighborhood. It concatenates these three representations into a single matrix and then applies PCA followed by Leiden clustering to obtain domain clusters.

**SpatialPCA** is a spatially aware dimension reduction method aimed at inferring a low-dimensional representation of the spatial transcriptome data^13^. In particular, SpatialPCA factorizes the gene expression matrix into a pair of matrices: the loading matrix and hidden factors. To reflect the spatial information, SpatialPCA assumes that each hidden factor is derived from a multivariate normal distribution function where its covariance matrix models the correlation among the spatial locations. Lastly, it uses Louvain clustering to determine the spatial domains.

**SCAN-IT** is a deep learning-based method for spatial domain identification^7^. It first builds a spatial network where nodes represent cells. Next, it uses a graph convolutional neural network to represent cells in an embedding space. Finally, it applies Leiden clustering on cells in the embedding space to identify spatial domains.

**BASS** employs a hierarchical Bayesian approach for spatial domain segmentation. It models three key probabilities: the likelihood of observing specific gene expression in a cell, conditional on its type; the likelihood that a cell belongs to a particular type, conditioned on its spatial domain; and the likelihood of a spatial domain as a function of the neighborhood graph. Using this model, the spatial domains are determined through Bayesian inference.

### Datasets and ground truth annotations

#### Dataset 1

This dataset was acquired via MERFISH technology^25^ and published in 2018. The number of measured genes is 155 and cells were segmented using a seeded watershed algorithm applied to the DAPI and total mRNA co-stains. Originally the dataset contains 12 samples of which only five were annotated with spatial domain labels^11^. These samples contain between 5,000 and 6,000 cells each. We obtained the annotated dataset from the resource provided by Yuan et al.^1^ (Supplementary Table 1).

#### Dataset 2

This dataset contains STARmap measurements for three mouse medial prefrontal cortex samples with 166 genes and one sample of the mouse visual cortex measured with Starmap technology and 1,020 genes^17^. Cell segmentation was performed by manual cell nuclei identification and a subsequent cell body segmentation based on a Nissl staining. The layers were previously annotated by an expert^11^. The four samples contain between 1,000 and 1,200 cells each. We downloaded the data from the resource provided by Yuan et al.^1^ (Supplementary Table 1).

#### Dataset 3

This dataset was measured using the BaristaSeq technology and contains three samples derived from the mouse cortex^16^. The number of measured genes is 79 and each sample contains between 1,500 and 2,000 cells. Spatial domains have been manually annotated^26^. We downloaded the data from the resource provided by Yuan et al.^1^ (Supplementary Table 1).

#### Dataset 4

This publicly available dataset contains three mouse brain samples measured with Xenium technology by the manufacturer 10x Genomics. The number of unique genes is 248 and cells were segmented by 10x Genomics using their proprietary cell segmentation pipeline. Each sample contains about 150,000 cells. The dataset has been recently annotated by Bhuva et al.^2^ using the Allen Brain Reference Atlas^27^. They performed a transcript-level annotation of spatial domains, which allowed us to easily consider two different cases per sample, one only using the transcripts measured inside the nuclei and another one using all transcripts measured inside the segmented cells (Extended Data Fig. 3). We removed all dissociated cells, by defining a cell graph based on a distance of 60 microns and removing all cells that were not part of the largest connected component. We downloaded the data from the resource provided by Bhuva et al.^2^.

#### Dataset 5

This publicly available dataset was measured with the MERFISH technology by the manufacturer Vizgen. The number of measured genes is 483 and cells were segmented by Vizgen with the default Vizgen segmentation pipeline. Since the dataset lacks ground truth domain annotations we developed an automated annotation workflow similar to the one by Bhuva et al. (see next section). Originally the dataset contains nine samples derived from the mouse brain of which we only consider the four most symmetric ones, to ensure the quality of our automated domain annotation workflow. The four samples each contain about 80,000 cells. The links to the raw data can be found in Supplementary Table 1.

#### Dataset 6

This dataset contains 31 mouse brain samples measured with MERFISH technology^19^. The number of unique genes is 374 and cells were segmented by the authors using Cellpose^28^. The spatial domains were annotated based on the manual cell type annotations. First, they overclustered each cell’s neighborhood cell type composition after PCA and then manually merged subclusters to obtain eight domain clusters. We downloaded the data from CELLxGENE using the link provided in Supplementary Table 1.

We followed the same preprocessing strategy for all datasets, namely removing all cells with less than 10 counts and removing all genes expressed in less than 5 cells. A detailed overview of all datasets and their sources can be found in Supplementary Tables 1 and 2.

### Automated annotation workflow

To annotate the ground truth spatial domains for Dataset 5, we used the Allen Brain Reference Atlas^27^, which provides a 3D reference volume of an entire mouse brain. First, an axial slice that best matches the anatomical regions of the DAPI image was selected manually via the Allen Python SDK for every individual MERFISH slice. Next, the reference slices and DAPI images were downscaled to a resolution of 25 microns per pixel to reduce computational cost. We then used the Elastix^29^ toolkit and the pyelastix Python wrapper to register the images in a two-step process. First, a rigid transformation was computed that allows only translational and rotational transformation of the moving image. Subsequently, an affine transformation was applied that allows shearing and stretching. The second step is particularly important since the orientations of the utilized MERFISH slices did not always perfectly match the axial view of the reference volume and were not perfectly symmetric. Mutual information was used as a similarity metric between the two images and stochastic gradient descent-based optimization was performed until convergence. Finally, the annotated axial slice corresponding to the DAPI image was transformed according to the previously computed parameters, and MERFISH cell coordinates were assigned based on the overlap of their respective centroids with a specific annotation ID.

### Evaluation metrics

To evaluate and compare the performances of the different spatial domain identification methods in recognizing the “ground-truth” labels (we also refer to as class), we considered the following evaluation metrics:

#### Normalized mutual information score (NMI)

NMI is a measure used to evaluate the similarity between two data clusterings (i.e. the model-based and ground-truth) based on their mutual information. The value is normalized between 0 and 1. A score of 1 indicates perfect clustering correspondence, while a score of 0 indicates no clustering correspondence. We implemented NMI via the *normalized_mutual_info_score* function from the scitkit-learn (v1.3.0) Python package.

#### Homogeneity score (HOM)

HOM is a measure indicating the purity of clusters with respect to the ground-truth labels. It quantifies whether each cluster predominantly consists of data points from a single class. A HOM of 1 indicates perfect homogeneity, while a score of 0 indicates poor homogeneity. We implemented HOM via the *homogeneity_score* function from the scitkit-learn (v1.3.0) Python package.

#### Completeness score (COM)

COM is a measure to evaluate the extent to which all members of a given class are assigned to the same cluster recognized by the models. A COM of 1 indicates that the model perfectly clusters all members of each class, while a score of 0 suggests that members of the same class are scattered across different clusters. We implemented COM via the *completeness_score* function from the scitkit-learn (v1.3.0) Python package.

### Benchmarking workflow and setup

On Datasets 1 to 3, we reused the quantitative results provided by Yuan et al. and compared them against NichePCA, Banksy, and SpatialPCA using a similar benchmarking workflow. In particular, the number of domain clusters identified with each method was chosen to fit the ground truth annotations for each dataset. Additionally, for NichePCA and Banksy we varied the number of nearest neighbors in the range from 5 to 20 and chose the value with the best performance across all datasets and all three metrics. For SpatialPCA we tested different kernel bandwidths in the range from 0.1 to 0.5, again selecting the value showing the best performance. To produce a visual plot of the domains identified by SCAN-IT for a selected sample from Dataset 1 we used the same settings as Yuan et al.

On Datasets 4 and 5, we only tested the top three performing methods besides NichePCA: Banksy, SCAN-IT, and SpaceFlow. We were not able to run BASS or SpatialPCA within a memory budget of 128 GB. For each method and sample of Dataset 4 and 5, we varied the number of nearest neighbors for spatial graph construction between 5 and 29 and the resolution for Leiden clustering (used by all four methods by default) between 0.1 and 2.0 with a step size of 0.1 while restricting the maximum number of clusters to 60. Then we selected the best-performing resolution across the average of all metrics for each method, each sample, and each number of the nearest neighbors setting. Lastly, we selected the number of nearest neighbors per method and dataset (not sample) such that the average performance across all metrics and the previously determined resolution settings is maximal. The corresponding metrics for these runs were then compared between methods. Since for Dataset 5, some of the cells could not be assigned to any of the atlas reference domains we ignored them during metric calculation.

### Multi-sample integration

To obtain coherent spatial clusters across multiple samples, we utilized the Python implementation of Harmony accessible via Scanpy (v1.9.8). Importantly, the graph construction and feature aggregation were performed separately for each sample while the PCA was applied across all samples. Finally, we performed Leiden clustering to obtain the spatial domains. We applied this workflow to Dataset 6 and considered the 26 nearest neighbors for graph construction. We selected a Leiden resolution of 0.15, which produced a similar number of clusters per sample as in the ground truth annotations. For visualization, we matched the cluster colors with their corresponding domains in the ground truth annotations (Fig. 2b).

### Scalability analysis

We measured compute times and memory usage for NichePCA, Banksy, SpaceFlow, and SCAN-IT on multiple subsampled versions of a selected sample of Dataset 4 with cell numbers ranging from 5,000 to 155,000. We executed each method 10 times per subsample and chose the median value for both measures. We also tracked the clustering time for SCAN-IT, SpaceFlow, and NichePCA, as these methods use a separate Leiden clustering step. All methods were executed in a single thread on an AMD EPYC 7742 64-core processor.

### Interpretability

On MERFISH brain data, we measured the variation explained by the principal components (PCs) and assessed the feature loadings of the first PC using Scanpy (v1.9.8). The PC coordinates of the 54 cluster centroids were computed by taking the mean of the 30 principal components, resulting in a 54 × 30 matrix. This matrix was multiplied with the PCA feature loadings (a matrix of dimension 30 × 483), resulting in a 54 × 483 matrix with gene contributions to each cluster.

### Ablation study

We conducted an ablation study to understand better which components of our computational pipeline are critical for identifying spatial domains. Therefore we considered all possible variations involving the same or fewer steps than the original NichePCA, namely median normalization, log-transformation, mean neighborhood aggregation, and PCA. Since we can presume that the aggregation step is critical we only considered one algorithm without it. For comparability, we applied all ablations and NichePCA on Datasets 1-3 using the same number of nearest neighbors (K=7).

### Computational resources

All analyses were performed on a server with an AMD EPYC 7742 central processing unit (2.25 GHz, 512 MB L3 cache, 64 CPU cores in total), and 128 GB of memory.

## Data availability

All the raw data can be downloaded from the links provided in Supplementary Table 1. The workflows for further processing and domain annotation are described in the Methods section.

## Code availability

NichePCA is available as a Python package at https://github.com/imsb-uke/nichepca. All scripts to reproduce the benchmarking and analyses will be made available after publication at https://github.com/imsb-uke/nichepca-analysis.

## Acknowledgments

We would like to thank the members of the Institute of Medical Systems Biology for their feedback and Sven Heins and Vadim Ustinov for IT support. This study was supported by grants from the Deutsche Forschungsgemeinschaft (DFG) to UP (SFB 1192 A1 and C3), CFK (SFB 1192 A5 and C3; KR 3483/3-1) and SB (SFB 1192 A2, B8, and C3). RK was supported by the 3R (Replace, Reduce, Refine) funding of the UKE. BY was supported by the Hamburg Macht Kinder Gesund foundation.

## Author contributions

Conceptualization: D.P.S., B.Y., and S.B. Methodology: D.P.S, B.Y., Formal analysis: D.P.S. Writing original draft: D.S.P., B.Y., N.K., and R.K. Writing review and editing: D.S.P., B.Y., N.K., R.K., and S.B. Visualization: D.S.P., B.Y., and R.K. Supervision: V.G.P., C.F.K., U.P., and S.B. Funding acquisition: C.F.K., U.P., and S.B.

## Figure legends

**Extended Data Figure 1:**
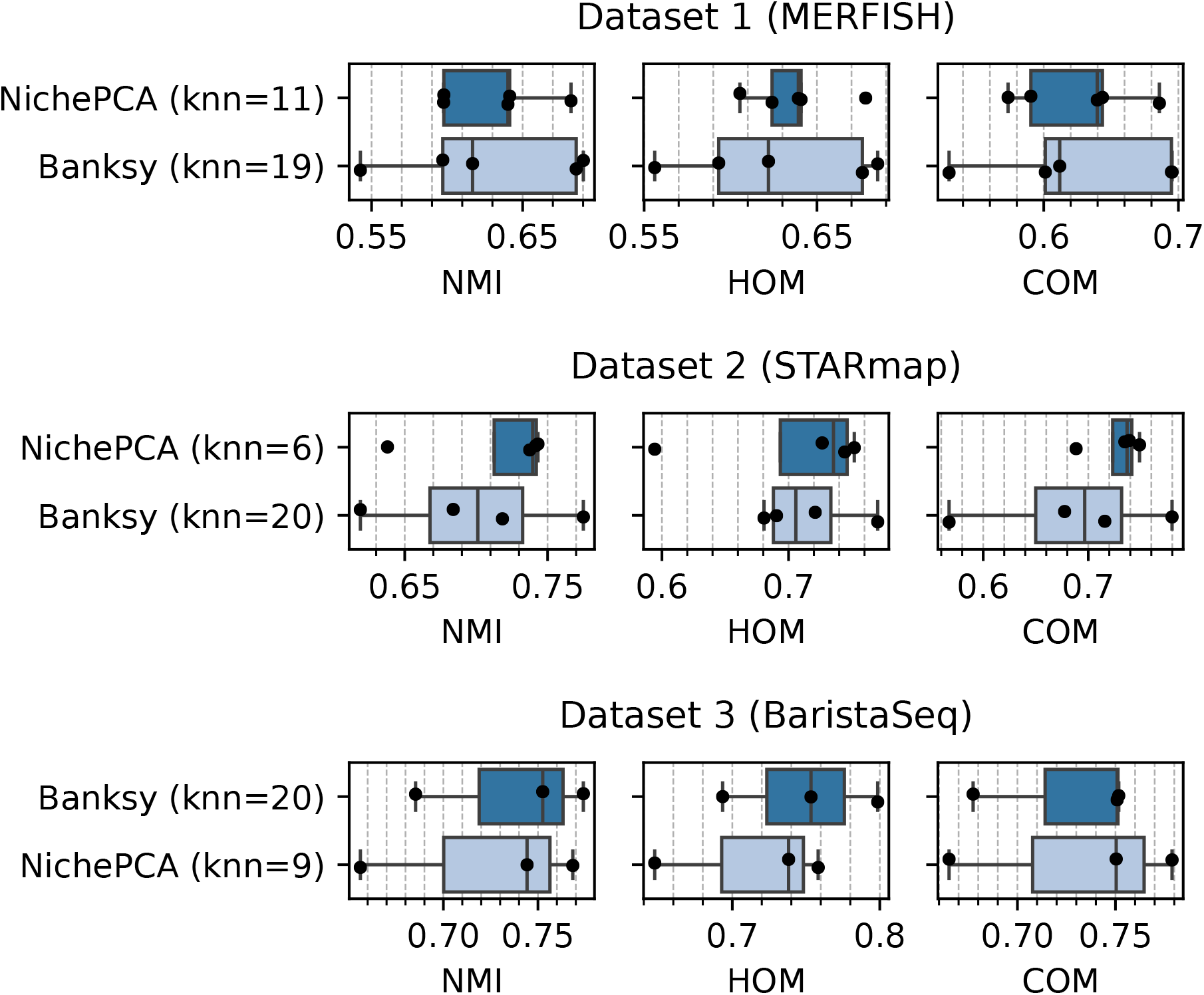
Performance in terms of NMI, HOM, and COM scores after optimizing the number of nearest neighbors per dataset for Banksy and NichePCA on Datasets 1-3 (top to bottom). The dots indicate the performance per sample. NichePCA outperforms Banksy in median NMI, HOM, and COM and features reduced performance variation.

**Extended Data Figure 2:**
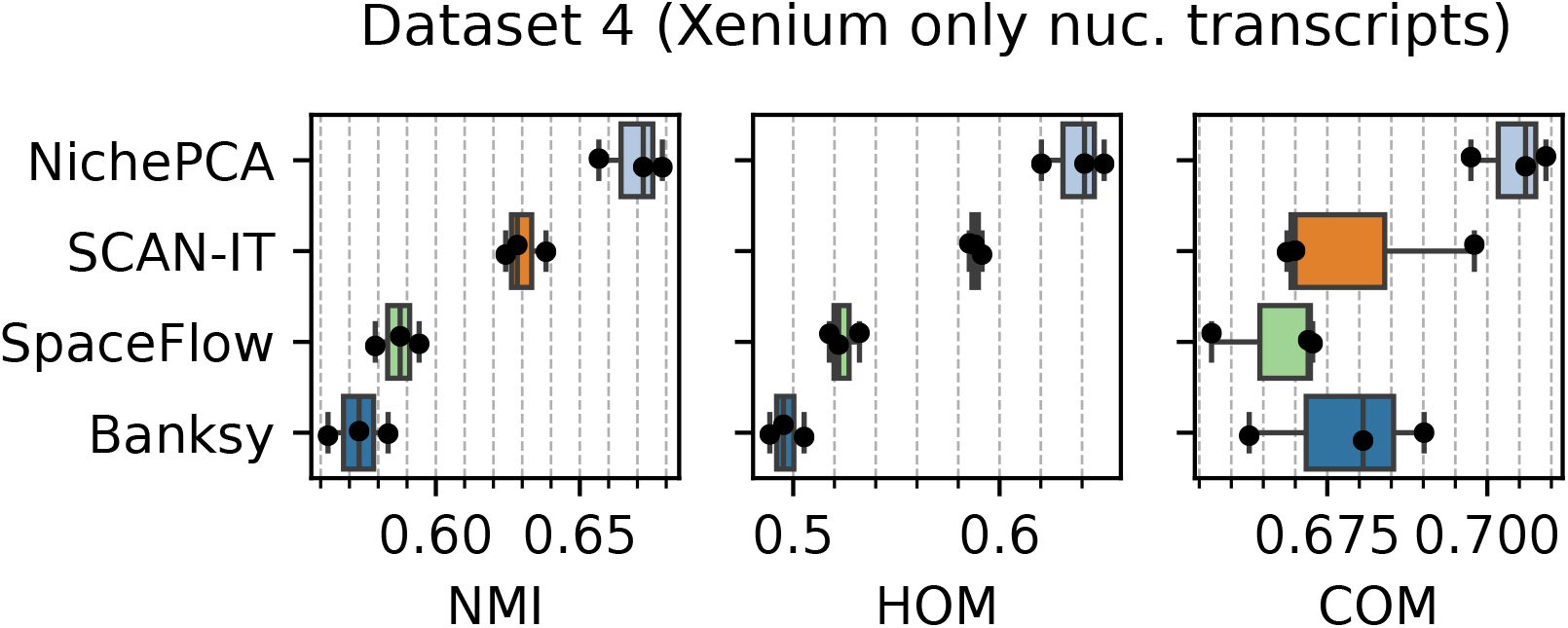
Ablation results for different variants of NichePCA on Datasets 1-3 regarding NMI, HOM, and COM scores. The annotations represent the steps of the different variants as follows: “agg” - mean neighborhood aggregation, “norm” - normalization to sum to the median value per feature, “log1p” - log-transformation with pseudocount of 1, “pca” - principal component analysis. Aggregation is always performed on a KNN graph with K=7. Dots indicate the performance per sample.

**Extended Data Figure 3:**
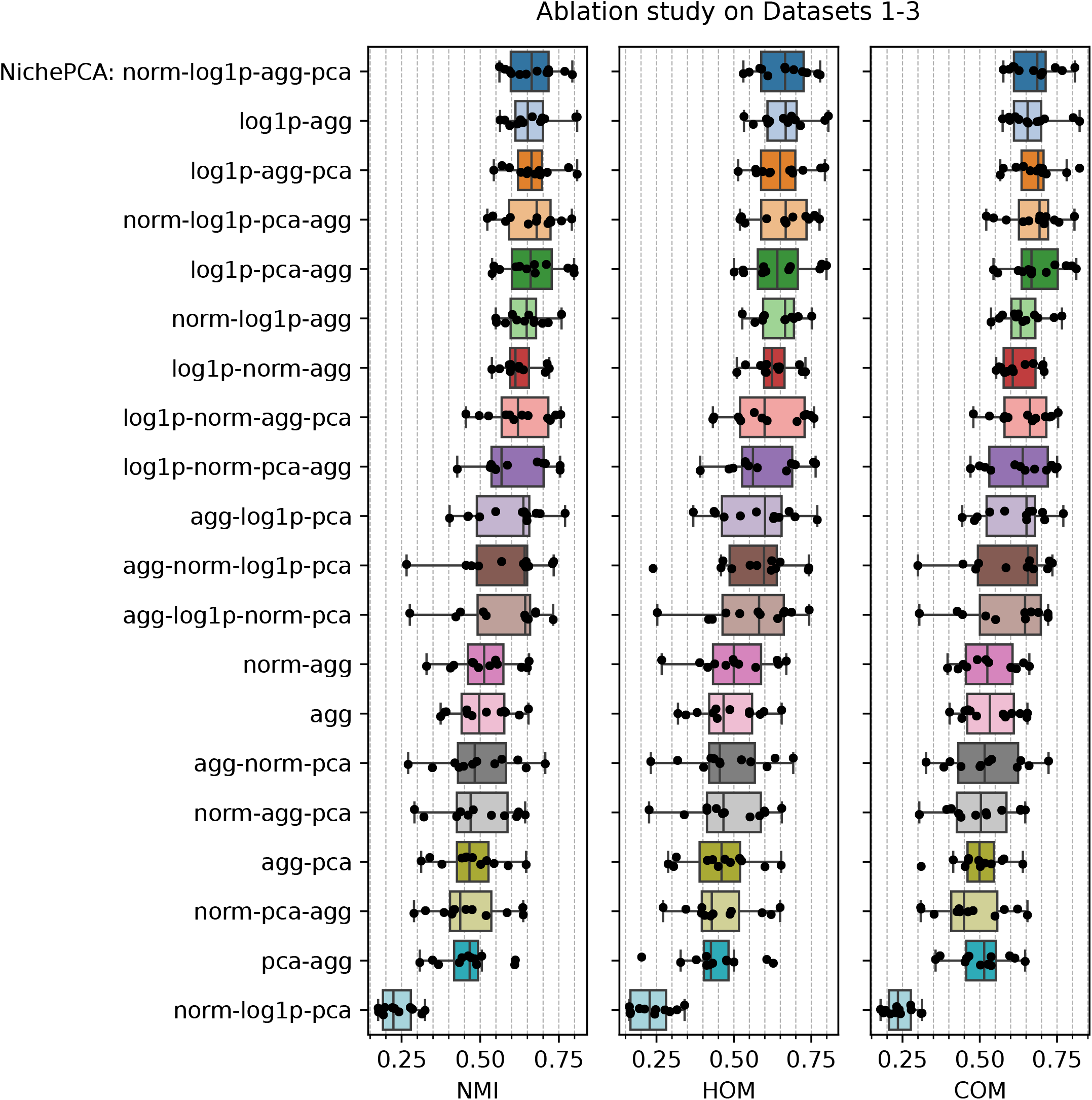
Performance in terms of NMI, HOM, and COM scores (left to right) for NichePCA, Banksy, SCAN-IT, and SpaceFlow on Dataset 4 (Xenium), considering only transcripts located within the segmented nuclei. The dots indicate the performance per sample.

**Extended Data Figure 4:**
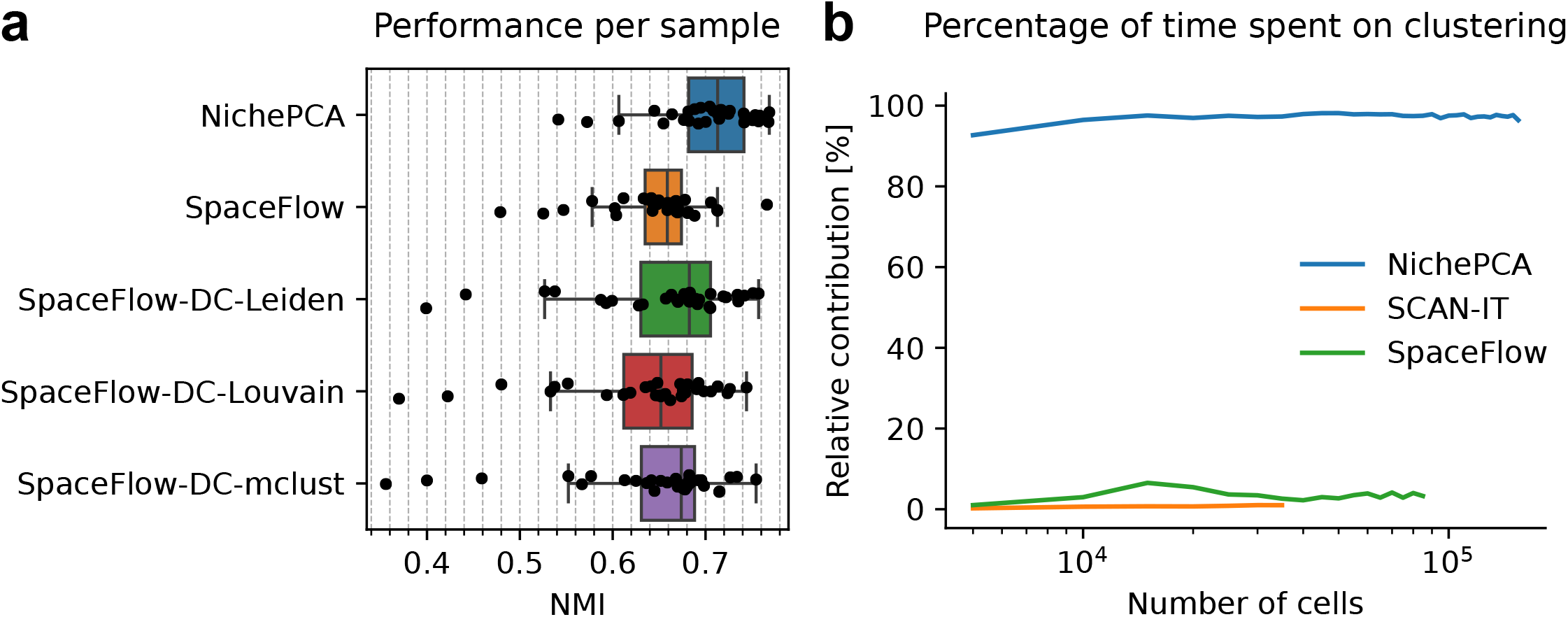
**a**, Spatial domain identification performance for all individual samples of Dataset 6 in terms of NMI score. Each dot represents the performance on one sample. **b**, Percentage of the total running time spent on clustering for NichePCA, SCAN-IT, and SpaceFlow. This evaluation is performed on the same sample of Dataset 4 and its subsampled versions as in Fig. 2c-d.

**Extended Data Figure 5:**
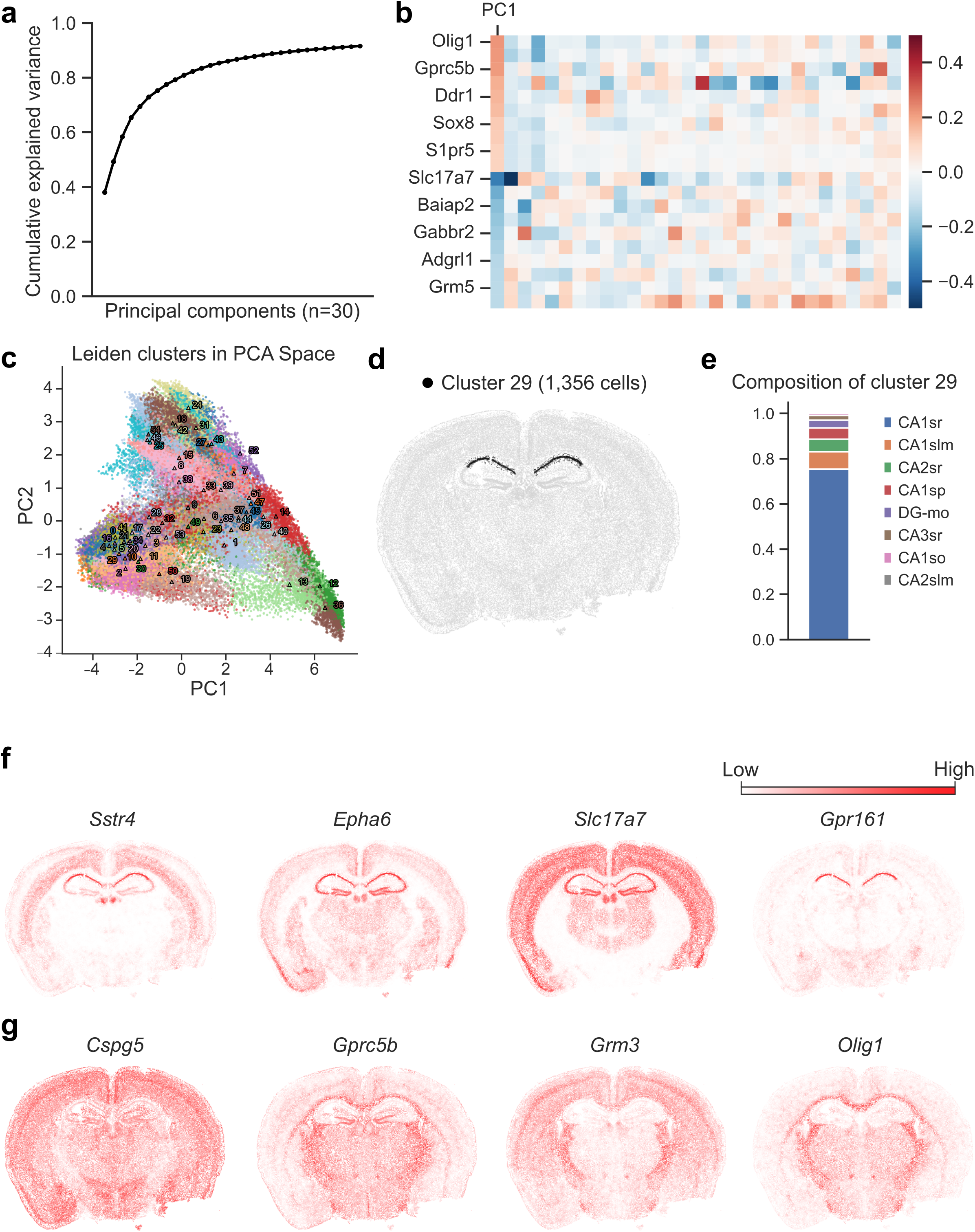
**a**, Cumulative variance explained by each principal component for a selected sample of Dataset 5. **b**, Loadings of genes with the most contribution to the first principal components. The five most positive and the five most negative genes are shown. **c**, Scatter plot showing the first and second principal components. The cells are colored by their cluster and the centers are labeled. **d**, Spatial localization of cells in cluster 29. **e**, Barplot showing the proportion of the ground truth domains within cluster 29. **f, g**, Expression patterns of genes with the most positive (f) and negative (g) contributions to the centroid of cluster 29.

